# Multimerin1, not Galectin-8, Promotes Gastric Chief Cell Differentiation by Tempering WNT Signaling

**DOI:** 10.1101/2025.11.04.685347

**Authors:** Xiaobo Lin, Gabriel Nicolazzi, Xuemei Liu, Chinye Nwokolo, Yeheil Zick, José B. Sáenz, Jeffrey W. Brown

## Abstract

Galectins are a family of proteins that bind galactose-containing glycans. One member, galectin-8, preferentially binds galactose that contains a terminal sulfate. Aberrant expression and secretion of sulfated glycosylation epitopes, such as 3’-Sulfo-Le^A/C^, is a feature of high-risk human foregut metaplasias. In addition, recent work has demonstrated that 3’-Sulfo-Le^C^ is a marker of mature murine zymogenic chief cells of the stomach and that 3’-Sulfo-Le^C^ epitope is secreted via cathartocytosis during the cellular transition to a metaplastic state. Based on those findings, we used *Lgals8^−/−^*mice, to determine whether galectin-8 might play a role in chief cell homeostasis. We observed delayed gastric differentiation in the *Lgals8^−/−^*mice and discovered that this phenotype was due to an unappreciated deletion of *Mmrn1* and *Snca* in the *Lgals8^−/−^* line. We show that multimerin-1 tempers WNT stimulation of the gastric corpus at an early age, as evidenced by nuclear beta-catenin staining and proliferation throughout the gland. Because multimerin-1 is synthesized and secreted from endothelial cells and not from the epithelial compartment, these data uncover a role for mesodermal cells in epithelial developmental and maturation of the mouse stomach. As prior studies have suggested galectin-8 and multimerin-1 have overlapping functions albeit, divergent with respect to bone, future studies using pure knockouts are necessary to refine these phenotypes.

## Introduction

The aberrant expression and secretion of sulfated glycosylation epitopes like 3’-Sulfo-Le^A/C^ is a feature of high-risk human foregut metaplasias, including Barrett’s esophagus, type III intestinal metaplasia of the stomach, and Pan-INs (pancreatic intraepithelial neoplasia), as well as in adenocarcinomas that arise from these metaplastic epithelia.^1–4^ The expression of sulfated glycotopes differentiates type III (highest risk; sulfated) from type II (moderate risk; sialylated) intestinal metaplasia of the human stomach. Concordant with the aberrant expression of these sulfated Lewis glycotopes in human metaplasias is the increased expression of the proteins that bind them (a subset of galectins).^5–9^

Galectins are a family of lectins with a common affinity towards galactose containing N-Acetyllactosamine moieties. This affinity is conferred by one or more carbohydrate recognition domains (CRD), which are a conserved fold formed by two antiparallel, 5-6 strand beta-sheets that lie against one another^10–14^. At least three of these galectins (galectin-3,^15^ galectin-4,^16^ and galectin-8^10,17,18^) have been shown to preferentially bind sulfated glycosylation epitopes relative to unsulfated moieties. Since both galectin-3^9,19,20^ and galectin-4^21–23^ have been demonstrated to play roles in foregut cancer biology and metastasis, we asked whether galectin-8 might also function along the gastric cancer cascade.

The mouse stomach differs from the human stomach in that sulfated mucins are expressed at homeostasis but secreted as the cells transition to metaplasia.^24^ In addition, there are several convenient, synchronous models to induce spasmolytic polypeptide-expressing metaplasia (SPEM) in the murine stomach^25–28^, a pre-neoplastic gastric lesion, which permits investigation into the specific roles proteins have on these sulfated mucins at homeostasis and following injury.^29,30^

By investigating alterations in chief cell maturation in the sole *Lgals8^−/−^* line,^31^ we discovered an unappreciated deletion of *Mmrn1* and *Snca* in *Lgals8^−/−^* mice that is responsible for the delay in chief cell maturation. Knowledge of this genomic deletion is important because galectin-8 and multimerin-1 have overlapping physiologic functions including regulating bone density^31–35^ and factor V coagulation.^36–40^ Mechanistically, we propose that multimerin-1 promotes gastric chief cell differentiation by tempering WNT stimulation during murine early gastric maturation.

## Results

### A Developmental Delay in Gastric Chief Cell Maturation

Unlike the human stomach, the zymogenic chief cells of the murine gastric corpus express sulfated mucins at homeostasis, which can be identified by the monoclonal antibody Das-1, which recognizes 3’-Sulfo-Le^C^ (Figure 1).^24,41^ Sulfomucin expression colocalizes with gastric intrinsic factor (GIF)^24^, a marker of differentiated chief cells, by ∼3 weeks of age in C57BL/6J mice and can serve as a marker of chief cell maturation (Figure 1C,D, Figure 2A). In *Lgals8^−/−^* mice, chief cell maturation was delayed on average by ∼25% postnatal life or ∼1 week (Figure 1G,H, Figure 2B,C). By 6 weeks of age, this significant histologic difference in sulfomucin or GIF expressing glands is lost (>200 corpus glands per mouse quantitated). However, we observe regions of consecutive glands where no mature chief cells are present at 6 weeks and later, which are not observed in wild-type mice (Figure 1E,F). Despite the gross histology phenotypes coalescing, we find that *Gif*, *Pgc*, and the chief cell scaling factor *Bhlha15*^42,43^ in *Lgals8^−/−^*mice do not normalize to age matched C57BL/6J (Figure 2D-F), suggesting that these maturation delays persist.

**Figure 1.**
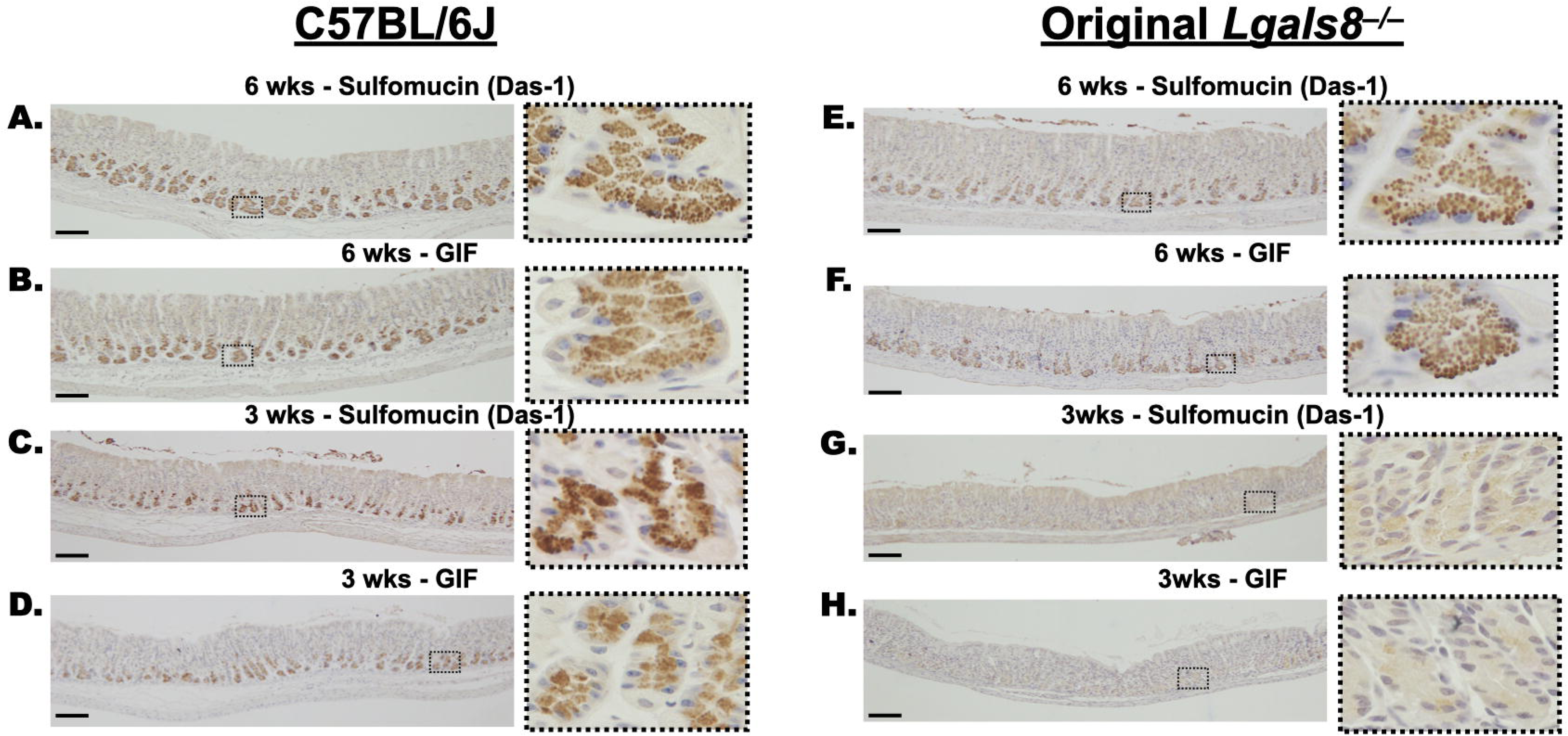
Delayed Maturation in the Original Galectin-8 Knockout Mouse. Mature, differentiated gastric corpus chief cells from wild-type C57BL/6J mice at 6 weeks of age express (**A**) sulfated mucins (recognized by the antibody Das-1) and (**B**) gastric intrinsic factor (GIF) in their zymogenic granules. **C,D.** Expression of sulfated mucins and GIF appear at 3 weeks of age in C57BL/6J mice. **E,F.** 6-week-old original *Lgals8^−/−^* mice express sulfated mucins and GIF in the chief cells, while **G, H.** 3-week-old original *Lgals8^−/−^*zymogenic granules as evidenced by the lack of Das-1 and anti-GIF reactivity. Scale Bars 100 μm.

**Figure 2.**
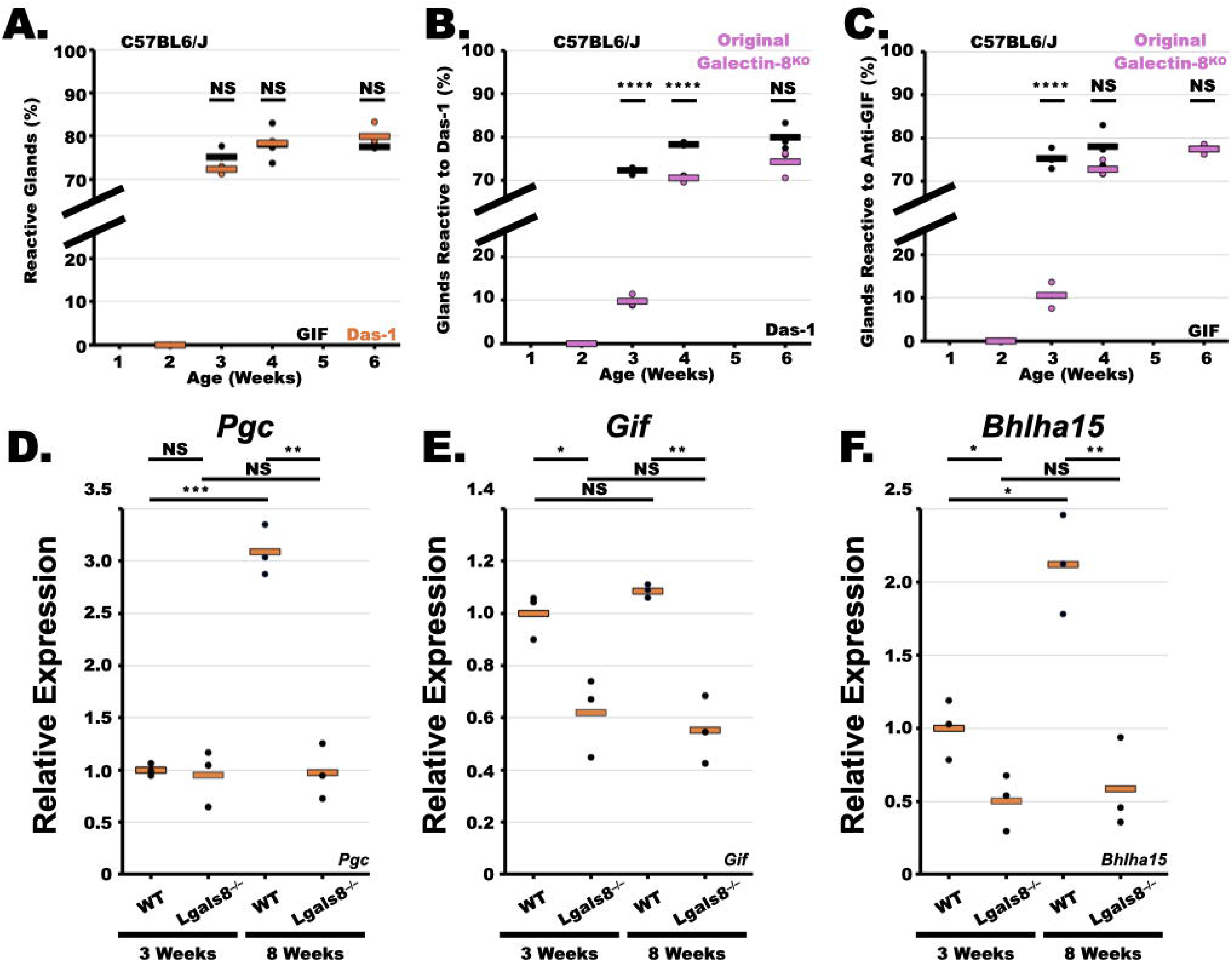
Quantification of Maturation Defect in the Original Galectin-8 Knockout Mouse. **A**. Sulfated mucins (orange) and gastric intrinsic factor (black) appear at equivalent ages in the gastric chief cell and serve as equivalent markers of chief cell maturation. The original galectin-8 null mice (purple) exhibit a delay in gastric chief cell maturation as measured by (**B**) Das-1 or (**C**) anti-GIF reactivity in comparison to C57BL/6 mice (black). Data points represent average of at least 200 glands per mouse and horizontal line represents average of three mice. Relative expression of **D.** pepsinogen C (*Pgc*) and **E**. gastric intrinsic factor (*Gif*), and **F.** Mist1 (*Bhlha15*) in the original *Lgals8^−/−^* line compared to C57BL/6J at 3 and 8 weeks of age. Black data points represent average signal from two reactions for each mouse. Orange bar is average of three mice. Significance calculated with ANOVA and post-hoc T-Test.

### Maturation Defect is not Observed in other Cell types in the Gastric Corpus

The gastric chief cell population is unique in that it is long-lived and self-sustaining^44,45^ cell type that does not depend on the isthmal transit amplifying population for its repletion, unlike other gastric corpus epithelial cell types (*e.g.* pit cells, neck cells, and parietal cells).^44,45^ Immunohistochemical analysis at three weeks of age with the lectins AAA (pit cells), GSII (neck cells), or ATP4B (parietal cells) did not demonstrate obvious defects in maturation, suggesting that the defect is specific to the chief cell compartment (Figure 3). We note a hazy, vesicular background in the AAA and GSII staining commonly seen in mature, chief cells in the wild-type C57BL/6J but not in the *Lgals8* knockout mouse, consistent with the developmental delay (Figures 3A,C).

**Figure 3.**
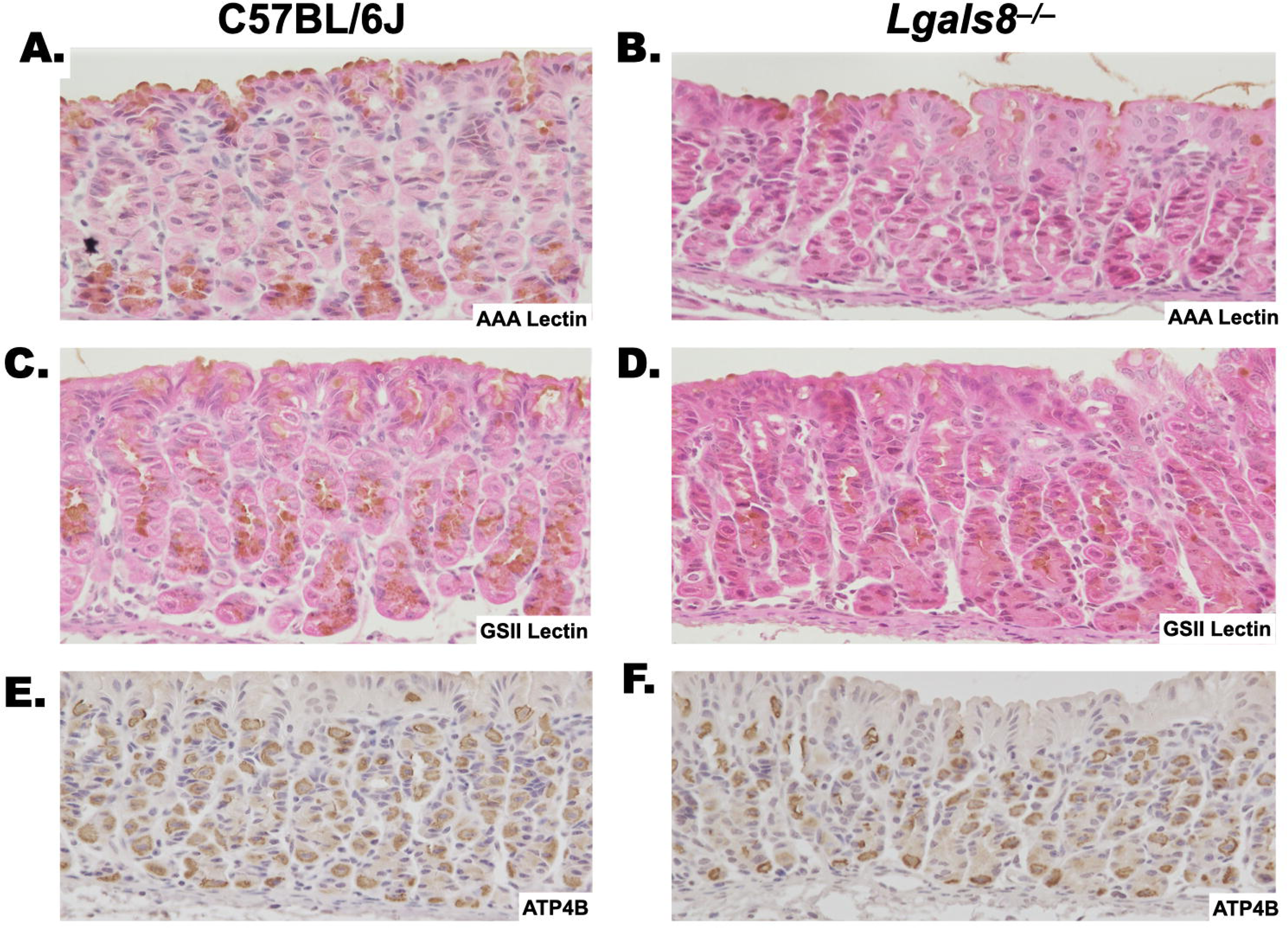
Cell types other than Chief Cells are Unaffected in the Original Galectin-8 null mouse. (**A**) Mature pit cells (AAA Lectin), (**B**) Neck cells (GSII lectin) and (**C**) Parietal cells (HK ATPase) are present at 3 weeks of age in both C57BL/6J and the original *Lgals8^−/−^* mice at 3 weeks of age.

### *Lgals8* is Not Expressed in the Zymogenic Chief Cells

Other than O-GlcNAc, all glycans are physically excluded from the cytoplasm. As such, at homeostasis, lectin-mediated phenotypes typically result from extracellular interactions with glycosylated membrane proteins as opposed to cell intrinsic / cell autonomous interactions. Using recently published single cell RNAseq dataset,^46^ we found that *Lgals8* is not expressed in the zymogenic chief cells, but instead in the tuft cells, pit cells, and proliferative cells (Figure 4B,C), which would suggest that galectin-8 might have a paracrine action on the zymogenic chief cell.

**Figure 4.**
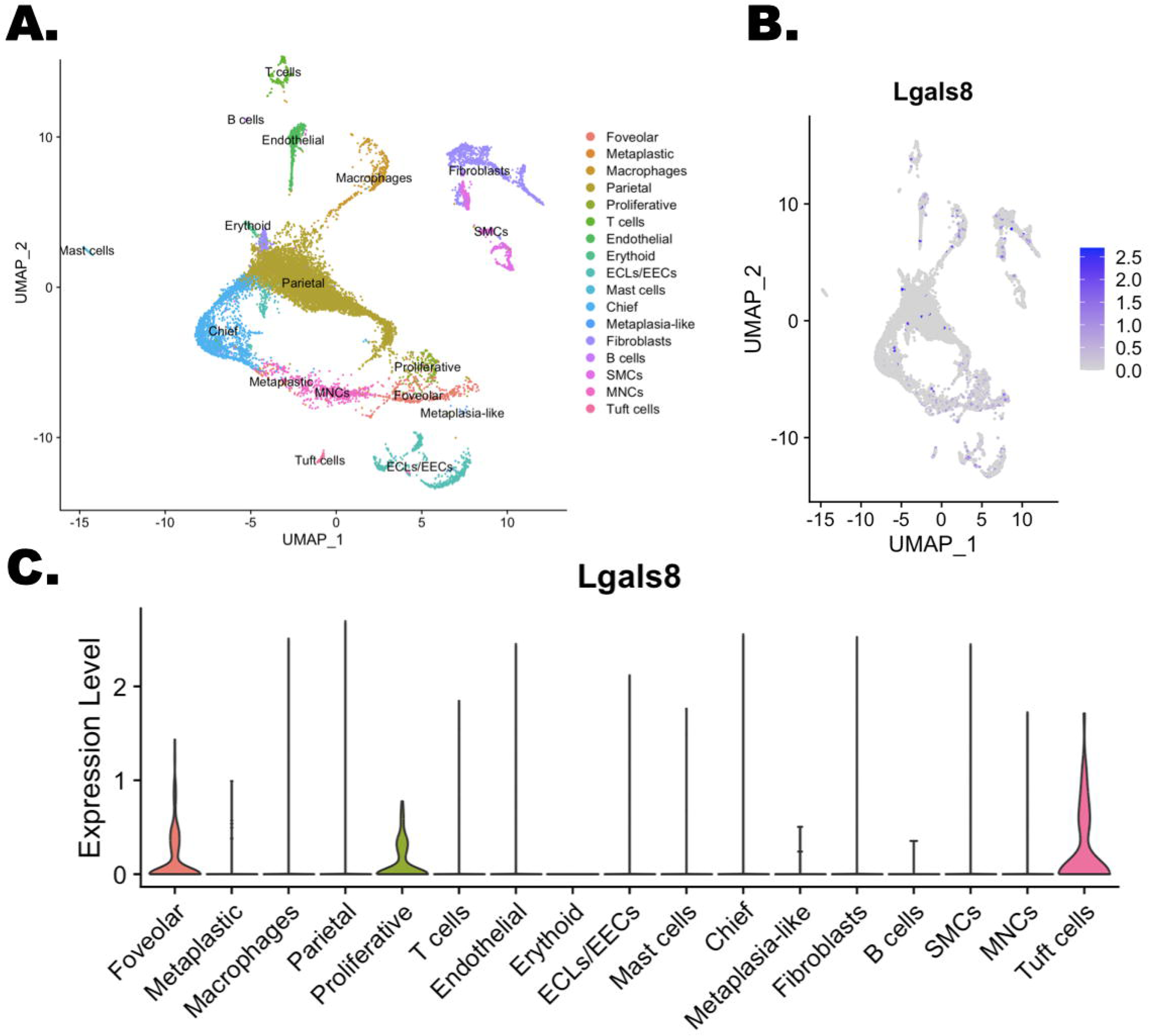
Single Cell RNA Sequencing Demonstrates Galectin-8 is not Expressed in Chief Cells. **A.** UMap of wild-type C57BL/6 mice at 6 weeks of age labeled by cell-type. **B**. Galectin-8 expressing cells overlaid on UMap. **C.** DIM plot displaying galectin-8 expression by cell-type.

Although we only observe phenotypic change in the maturation of zymogenic chief cells that represent a small subset of the cells in the gastric corpus, we proceeded with bulk RNAseq to determine transcriptional differences that might differ between the original *Lgals8^−/−^* mice and wild-type C57BL/6J. This was performed at 3 weeks of age, when there were significant differences in chief cell maturation, and at 8 weeks of age, when the gross histologic appearances are indistinguishable (Figure 2B,C). Despite the groups segregating from one another in principal component analysis (Figure 5A), few transcripts were significantly different between the groups (Supplemental Tables 1 and 2). One striking outlier between the groups was multimerin 1 (Supplemental Table 1), a large extracellular glycoprotein encoded by *Mmrn1* that has been studied in the contexts of coagulation cascade and bone density, two functions also affected by Galectin-8. As expected, bulk RNA sequencing revealed that *Lgals8^−/−^* mice were not only deficient in *Lgals8* but also in *Mmrn1*, which were confirmed qRT-PCR (Figure 5B,C). These findings suggested that the founder *Lgals8^−/−^* mice may exhibit unexpected features related to *Mmrn1* deficiency.

**Figure 5.**
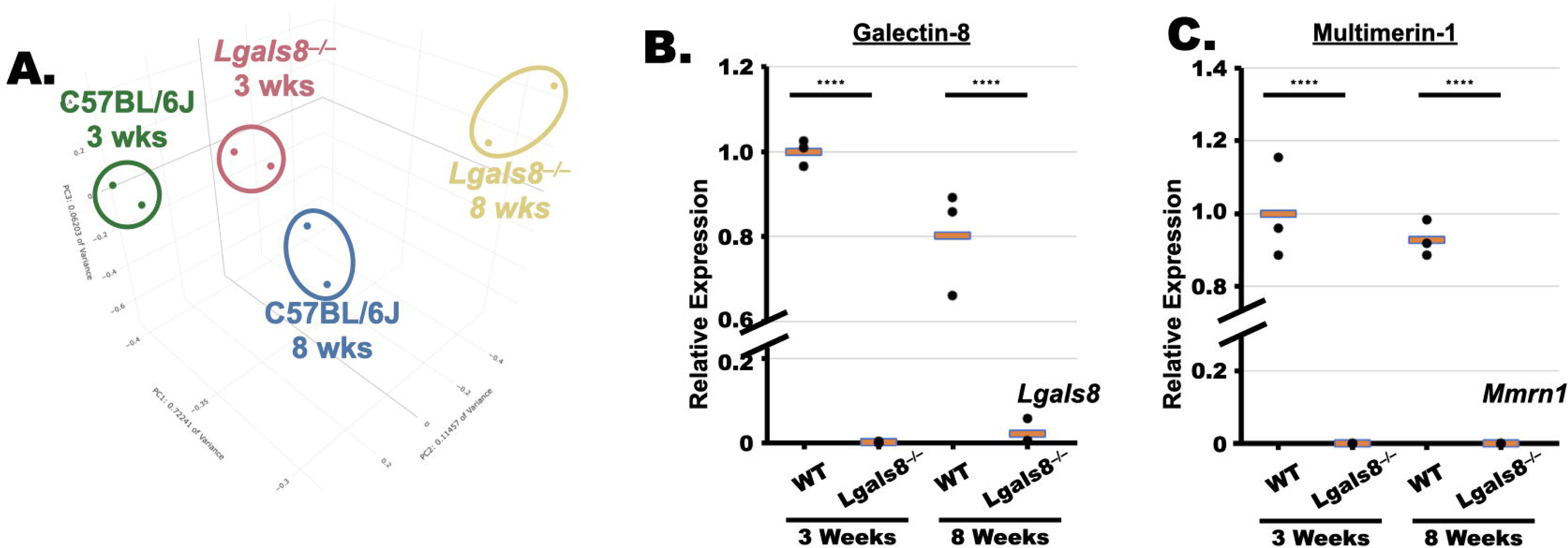
Bulk RNA sequencing Demonstrates Exceedingly Low Expression of Multimerin 1 in the Original Galectin-8 Knockout. **A.** Principal component analysis of C57BL/6J and the original galectin-8 null at 3 and 8 weeks of age demonstrating segregation of mice by genotype and age. **B.** qRT-PCR of *Lgals8* confirms knockout. **C.** qRT-PCR of *Mmrn1* confirms dramatic decrease in expression of Multimerin in galectin-8 null mice.

### Galectin-8 Knockdown Does not Affect Mmrn1 Levels

Multimerin1 is expressed in the endothelium as well as in platelets.^47^ To determine whether Galectin-8 alters the expression of *Mmrn1*, we knocked down *LGALS8* in HUVEC cells and assayed for *MMRN1* levels. Transfection of three different shRNA constructs against *LGALS8*, each resulting in ∼50% knockdown of *LGALS8* (Figure 6A) did not alter the levels of *MMRN1* (Figure 6B). These findings suggest that the loss of *Mmrn1* may not be a consequence of *Lgals8* deficiency.

**Figure 6.**
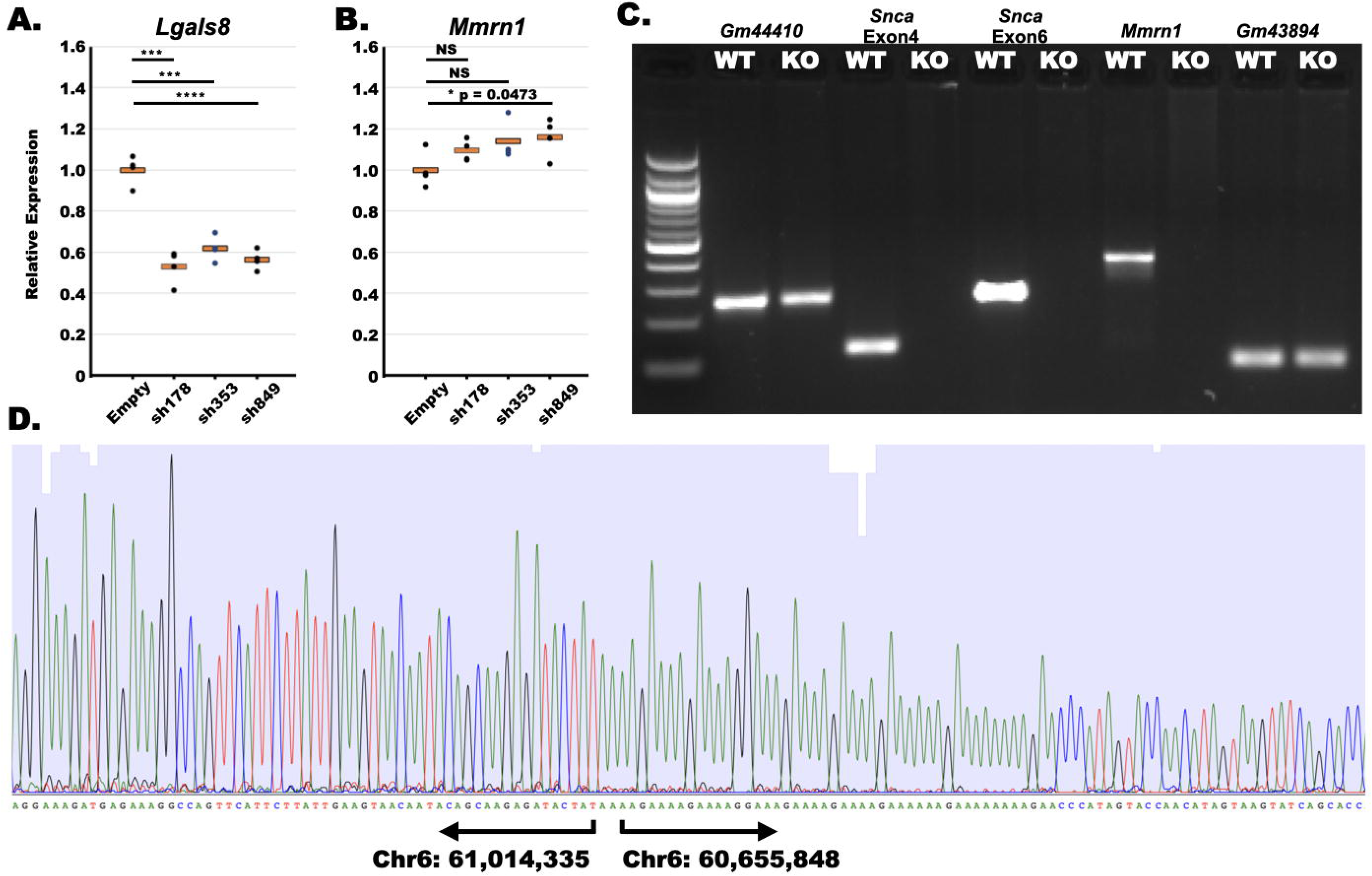
Unappreciated Genetic Deletion of Multimerin 1 in the Original Galectin-8 Null Line. shRNA against *Lgals8* reduces expression of (**A**) *Lgals8* but not (**B**) *Mmrn1* in HUVEC cells. **C**. PCR targeting alpha-synuclein, Multimerin-1 as well as upstream and downstream regions demonstrates genomic deletion of alpha-synuclein and Multimerin-1 in the galectin-8 null mice. **D**. Sequencing across the deletion reveals an identical deletion to that reported in the OlaHsd strain C57BL/6J.

### Galectin-8 Knockout Mice Contain a Deletion of Genetic Locus Containing Alpha-Synuclein and MMRN1

As the expression of *Mmrn1* was exceedingly low in the gastric corpus (Figure 5C) and knockdown of *LGALS8* did not alter the expression of *MMRN1* (Figure 6B), we genotyped the *Lgals8^−/−^* line for *Mmrn1* and discovered that *Lgals8^−/−^* mice carried an unknown deletion in the *Mmrn1* locus as well as in the neighboring alpha synuclein (*Snca*) locus (Figure 6C). The coordinates for the genomic deletion in the *Lgals8^−/−^* line were identical to the C57BL/6J-OlaHsd^48^ line. Those findings suggest that the *Lgals8^−/−^* contains a unique additional deletion that could be responsible for the delay in chief cell maturation.

### Multimerin-1 but not Galectin-8 is responsible for delayed chief cell maturation

Since murine *Lgals8* is located on chromosome 13, while *Mmrn1* & *Snca* are on chromosome 6, we outbred the *Lgals8^−/−^/Mmrn^−/−^1/Snca^−/−^*triple knockout mice onto congenic C57BL/6J mice to segregate the alleles. We found that chief cells in *Lgals8^−/−^* mice with wild-type Multimerin-1 and alpha-synuclein matured normally (Figure 7A-D), while *Mmrn1^−/−^/Snca^−/−^*mice, expressing wild-type levels of galectin-8, exhibited delayed chief cell maturation (Figure 8E-H). Consistent with alpha-synuclein being primarily expressed in the central nervous system, we could not detect mRNA for alpha-synuclein in the stomach (Supplemental Figure 1) with qRT-PCR, suggesting that that the delayed maturation phenotype is a consequence of loss of the endothelial glycoprotein multimerin-1.

**Figure 7.**
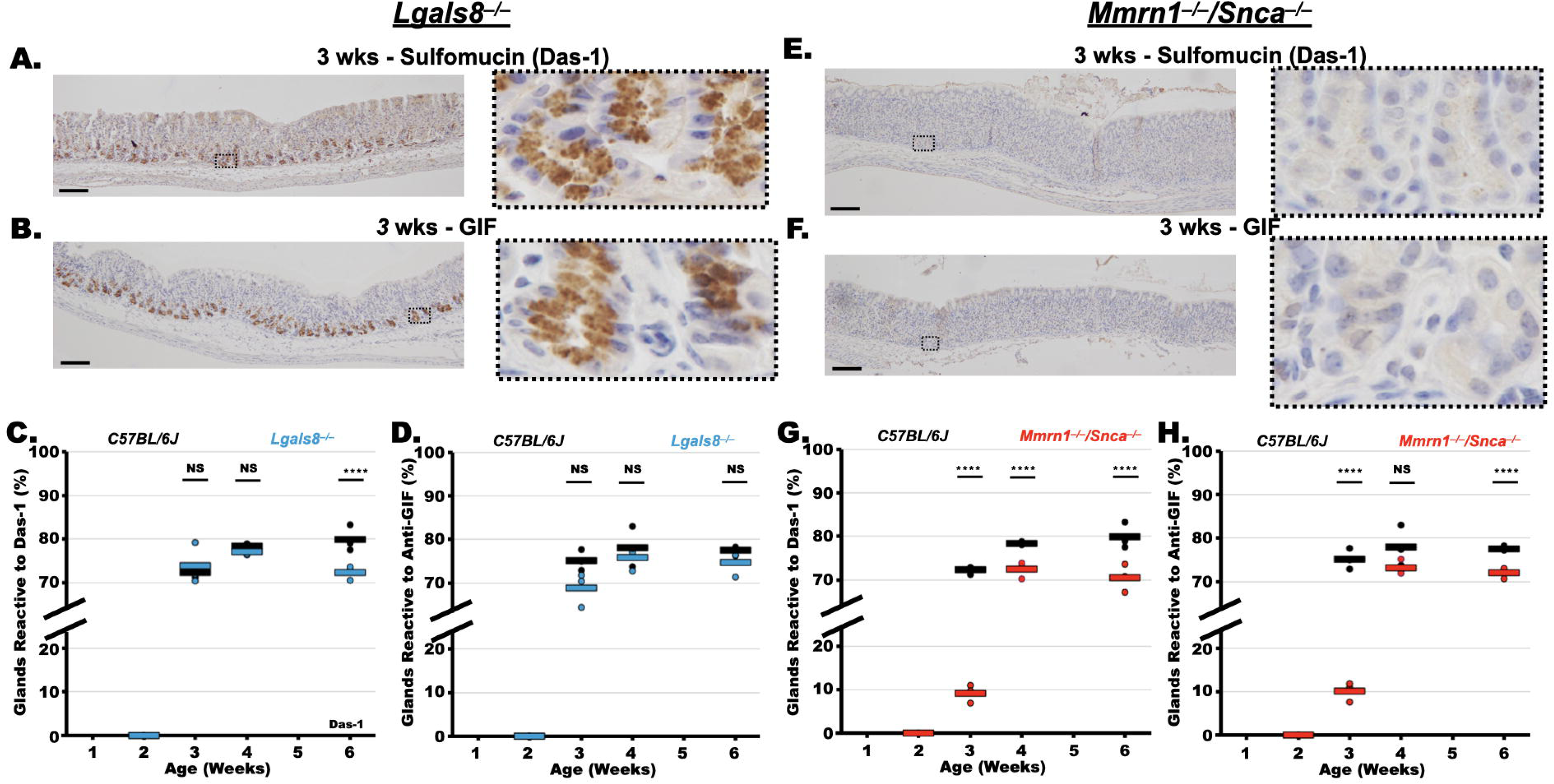
Delayed Chief Cell Maturation is a Consequence of Multimerin1 loss, Not Galectin-8. Immunohistochemistry of (**A**) sulfated mucins and (**B**) gastric intrinsic factor in pure galectin-8 mice is indistinguishable from wild-type C57BL/6J mice. Quantification of the percentage of corpus glands with either (**C**) sulfomucin or (**D**) gastric intrinsic factor expression in galectin-8 null mice compared to wild-type mice. Immunohistochemistry of (**E**) sulfated mucins and (**F**) gastric intrinsic factor in Multimerin-1/alpha-synuclein double knockout mice demonstrates delayed expression relative to wild-type C57BL/6J mice. Quantification of the percentage of corpus glands with either (**G**) sulfomucin or (**H**) gastric intrinsic factor expression in galectin-8 null mice compared to wild-type mice. Data points represent average of at least 200 glands per mouse and horizontal line represents average of three mice. Significance calculated with ANOVA and post-hoc T-test (statistical analysis needs refinement). Scale Bars 100 μm.

**Figure 8.**
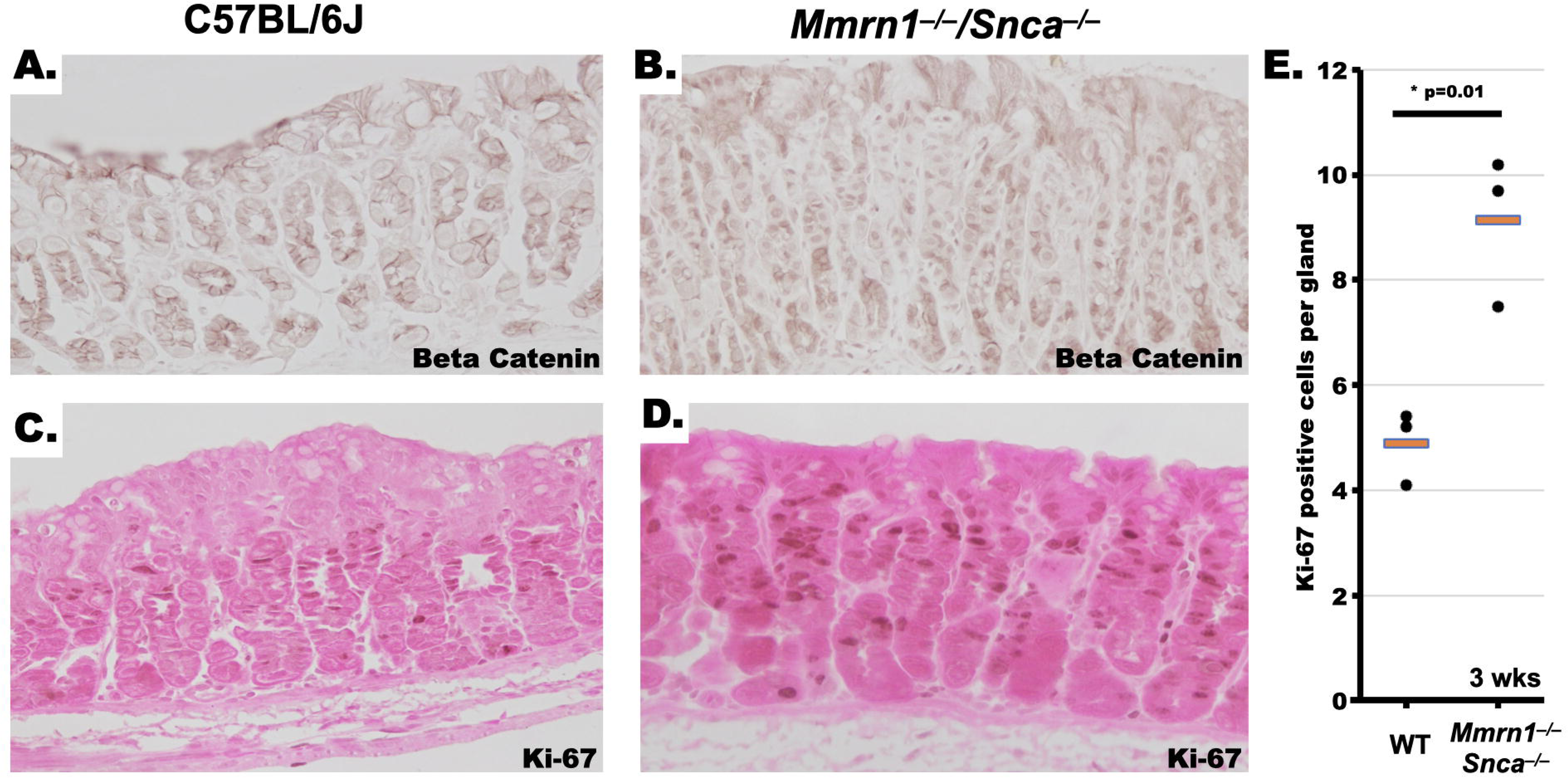
Multimerin-1 Loss Modulates Beta-Catenin Localization and Proliferation of the Gastric Corpus. **A.** Beta-catenin immunohistochemistry of wild-type C57BL/6J demonstrates membrane staining, while **B.** *Mmrn1^−/−^/Snca^−/−^* at 3 weeks of age displays prominent nuclear reactivity. **C,D.** Representative Ki-67 immunohistochemistry of wild-type C57BL/6J and *Mmrn1^−/−^/Snca^−/−^*, repectively at 3 weeks of age demonstrates greater proliferation in the *Mmrn1^−/−^ /Snca^−/−^* mice. **E.** Quantification of Ki-67 reactive nuclei per gland. Dots are average of >200 glands per mouse, orange line is the average of three mice. Significance calculated with T-Test.

### Multimerin-1 Tempers WNT Signaling in the Gastric Corpus

Homologs of multimerin-1 have been shown to bind and downregulate WNT through their EMI domain.^49,50^ To determine whether multimerin-1 acts similarly, we compared beta-catenin localization in wild-type mice to *Mmrn1^−/−^* stomachs at 3 weeks of age. In wild-type C57BL/6J mice, beta-catenin localized to the membrane, while in mice lacking *Mmrn1*, we observed a decrease in membrane staining and an increase in nuclear staining. Consistent with this staining pattern, we observed a greater number and wider distribution of Ki-67 positive cells in the stomachs of *Mmrn1^−/−^*mice, suggesting a proliferative phenotype. At older ages in the *Mmrn1^−/−^,* beta-catenin assumed the traditional membrane localization, suggesting that WNT signaling might be tempered by other redundant proteins as the mouse ages (Supplemental Figure 2).

## Discussion

Here, we attempted to explore the role of galectin-8 in the gastric epithelium during murine development. However, our investigation led us to discover an unappreciated deletion of multimerin 1 (and alpha-synuclein) in the galectin-8 line that is responsible for a maturation delay of the chief cell compartment of the gastric corpus observed in the original *Lgals8^−/−^* line.

Multimerin-1, a member of the EMILIN/multimerin family of proteins, is a large, glycoprotein secreted from the endothelium^47^ and megakaryocytes / platelets.^51^ Multimerin-1 has established roles is in thrombosis, where it binds activated platelets, von Willebrand factor (vWF),^52^ and factor V^40,53^, and extracellular matrix collagen^38^ to bolster platelet adhesive functions. In humans, secondary Multimerin-1 deficiency, resulting from excessive urokinase plasminogen activator in Quebec platelet disorder contributes to the disease phenotype.^54–57^ In addition to pro-thrombotic functions, deletion of multimerin-1 (and alpha-synuclein) in the C57BL/6J-OlaHsD mouse demonstrates bone loss.^32^

Galectin-8 has been shown to act in similar physiologic systems to multimerin-1. For example, overexpression of *LGALS8* causes increased secretion of RANKL resulting in increased osteoclastogenesis and ensuant bone mass reduction.^31,33,34^ The increased bone mass observed in the *Lgals8^−/−^* mouse was potentially abrogated by simultaneous deletion of multimerin-1, which has the effect of decreasing bone mass.^31^ Galectin-8 has been shown to activate platelets^58^ as well as bind factor V and function in endocytosis and storage factor V in platelet alpha granules,^36,59^ the same subcellular localization where Mmrn1 is stored. As such, any future studies investigating the role of galectin-8 using the murine knockout should only occur after outbreeding to wild-type mice with native multimerin-1 (and alpha-synuclein). We also found that in endothelial cells, knockdown of galectin-8 does not affect multimerin-1 levels suggesting independent roles (Figure 6).

Here, we uncovered a new function for multimerin-1 and a novel role of the endothelium promoting gastric maturation. We found that like other members of the EMILIN/Multimerin family,^49,50,60–63^ multimerin-1 affected WNT signaling. We found that beta-catenin was localized to the membrane in wild-type C57BL/6J mice at 3 weeks of age, but that this membrane localization was dramatically decreased with concurrent nuclear localization in the multimerin-1 null mice at 3 weeks of age. At older ages, beta-catenin staining assumed the default membrane localization in the multimerin-1. Due to the temporal association between beta-catenin localization, proliferative changes, and delayed chief cell maturation at three weeks of age, we propose that the delayed maturation phenotype in the multimerin-1 null mice is due to increased WNT stimulation. Multimerin-1 is an extracellular secreted glycoprotein that resides in the basement membrane and may act as a buffer for WNT signaling. In its absence, WNT hyperstimulation occurs.

The results presented here serve as (1) a cautionary tale of potentially attributing a phenotype to specific allele as whole genome sequencing is rarely performed for all murine genetic alleles (2) a novel role of the endothelium in promoting gastric maturation via secreting multimerin-1 which buffers WNT signaling.

## Methods

### Animals

All experiments using animals followed protocols approved by the Washington University in St. Louis, School of Medicine Institutional Animal Care and Use Committee. WT C57BL/6J mice were purchased from Jackson Laboratories (Bar Harbor, ME).

All mouse experiments were performed on mice aged 6-10 weeks and both sexes were utilized indiscriminately as prior studies have demonstrated an identical phenotype^26,27,64^.

Galectin-8 single knockout and Multimerin-1/Alpha-Synuclein double knockouts were generated by outbreeding the *Lgals8/Mmrn1/Snca* triple knockout to C57BL/6J mice Jackson Laboratories (Bar Harbor, ME) and identifying the alleles by genotyping. Pups were then backcrossed to generate the homozygous knockouts. They were maintained by breeding knockout to knockout.

### Imaging and tissue analysis

Following anesthetizing with isoflurane and cervical dislocation, murine stomachs were excised, flushed with PBS and fixed overnight with 10% formalin in PBS. They were washed and equilibrated in 70% ethanol for several hours prior to embedding in 3% agar and routine paraffin processing. Sections (5-7 μM) were prepared for immunohistochemistry and/or immunofluorescence by deparaffinization using Histoclear and an alcohol series for rehydration. Endogenous peroxidase was quenched by incubating slides in 1.5% hydrogen peroxide (Sigma, St. Louis, MO) solution in methanol for 15 minutes at room temperature. After that, antigen retrieval was performed in 10 mM citrate buffer, pH 6.0 in a pressure cooker. Tissue was blocked for 1 hour at room temperature with 2% BSA and 0.05% Triton X-100. Primary and secondary antibodies were diluted in 2% BSA and 0.05% Triton X-100. Vector ABC Elite kit was used for immunohistochemistry. Immunohistochemistry slides were mounted with Permount and immunofluorescence with Prolong Gold with DAPI. Brightfield images were taken on either a Nanozoomer (Hamamatsu 2.0-HT System) for quantitation or Olympus BX43 light microscope. Confocal images were obtained on a Zeiss LSM880 confocal microscope.

### Quantification and Statistical Analysis

Quantification of percent of Das1+ or GIF+ glands per mouse stomach was done in the same way. Each scanned image of IHC stained with Das1 or GIF was counted for Das1+ or GIF+ glands and total number of glands for three areas (approximately 25 glands for each area, totaling 75 glands per ring or 225 glands for 3 rings of each stomach). The percentage of Das1+ or GIF+ glands per mouse was calculated after dividing the sum of Das1+ or GIF+ glands by the grand total.

All statistical details can be found in the figure legends.

### Genotyping for Genomic DNA Deletion and Knockout of *Mmrn1/Snca*

Extracted mouse tail genomic DNA was used to examine the *Snca/Mmrn1* locus for deletion detection. Additional primers, GM43894 and GM44410, were designed for genes around the Snca locus. PCR was performed using DNA Engine (MJ Research) with the protocol: 1 cycle of denaturing at 94C for 2 minutes, 35 cycles each of denaturing at 94C for 30 seconds, annealing at 55C for 30 seconds, and polymerization at 72C for 30 seconds; and 1 cycle of polymerization at 72C for 1 minute.

A separate three-primer set was designed to distinguish wild type, heterozygous, and homozygous for Mmrn1. The PCR protocol was identical as above except the polymerization time was 20 seconds.

### Genomic DNA Amplification and Sequencing

To confirm the deletion of *Snca/Mmrn1* locus, primers were designed to amplify the genomic DNA around the deleted region by PCR. The PCR protocol was as follows: 1 cycel of denaturation at 94C for 2 minutes, 10 cycles each of denaturation at 94C for 20 seconds, annealing at 68C for 20 seconds with annealing temperature reduced 0.5C for each cycle, and amplifying at 72C for 50 seconds; and 28 additional cycles each of denaturation at 94C for 20 seconds, annealing at 60C for 20 seconds, extension at 72C for 50 seconds. Purified PCD product was sequenced by Sanger sequencing method (Genewiz).

### qRT RNA

RNA was isolated using RNeasy (Qiagen) per the manufacturer’s protocol. The integrity of the mRNA was verified with a BioTek Take3 spectrophotometer and electrophoresis on a 2% agarose gel. RNA was treated with DNase I using in-column digestion (Qiagen), and 1 μg of RNA was reverse-transcribed with SuperScript III (Invitrogen) following the manufacture’s protocol. Measurements of cDNA abundance were performed by qRT–PCR using Applied Biosystems QuantStudio 3 Real-time PCR system. Power SYBR Green master mix (Thermo Scientific) fluorescence was used to quantify the relative amplicon amounts of each gene. Primer sequences are located in the key resource table.

### Bulk RNA Sequencing

Total RNA integrity was determined using Agilent 4200 Tapestation. Library preparation was performed with 5ug of total RNA with a RIN score greater than 8.0. Ribosomal RNA was removed by poly-A selection using Oligo-dT beads (mRNA Direct kit, Life Technologies). mRNA was then fragmented in reverse transcriptase buffer and heating to 94 degrees for 8 minutes. mRNA was reverse transcribed to yield cDNA using SuperScript III RT enzyme (Life Technologies, per manufacturer’s instructions) and random hexamers. A second strand reaction was performed to yield ds-cDNA. cDNA was blunt ended, had an A base added to the 3’ ends, and then had Illumina sequencing adapters ligated to the ends. Ligated fragments were then amplified for 16 cycles using primers incorporating unique dual index tags. Fragments were sequenced on an Illumina NovaSeq 6000 using paired end reads extending 150 bases.

### DIM Plot

## STAR METHODS

### KEY RESOURCE TABLE

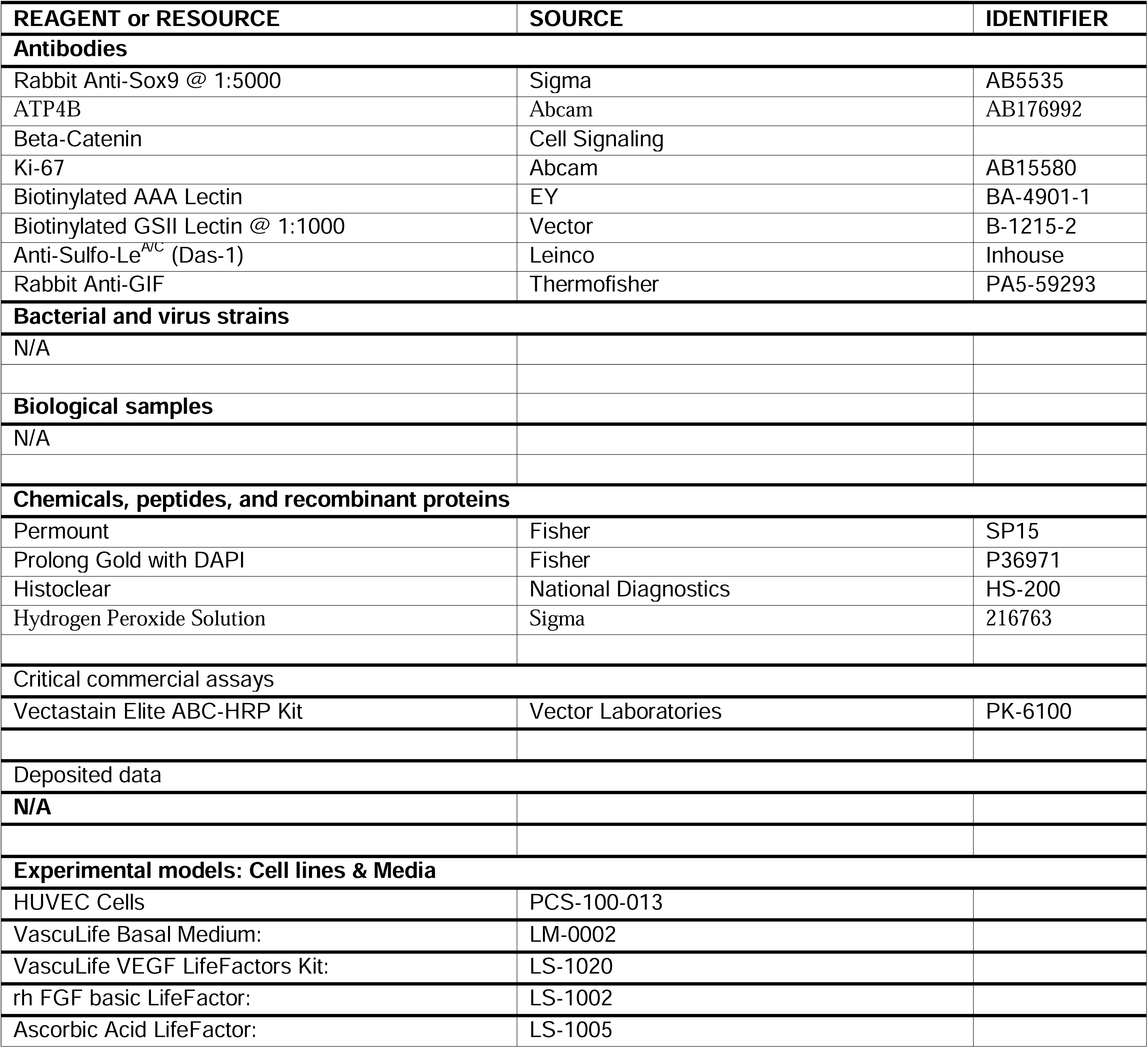
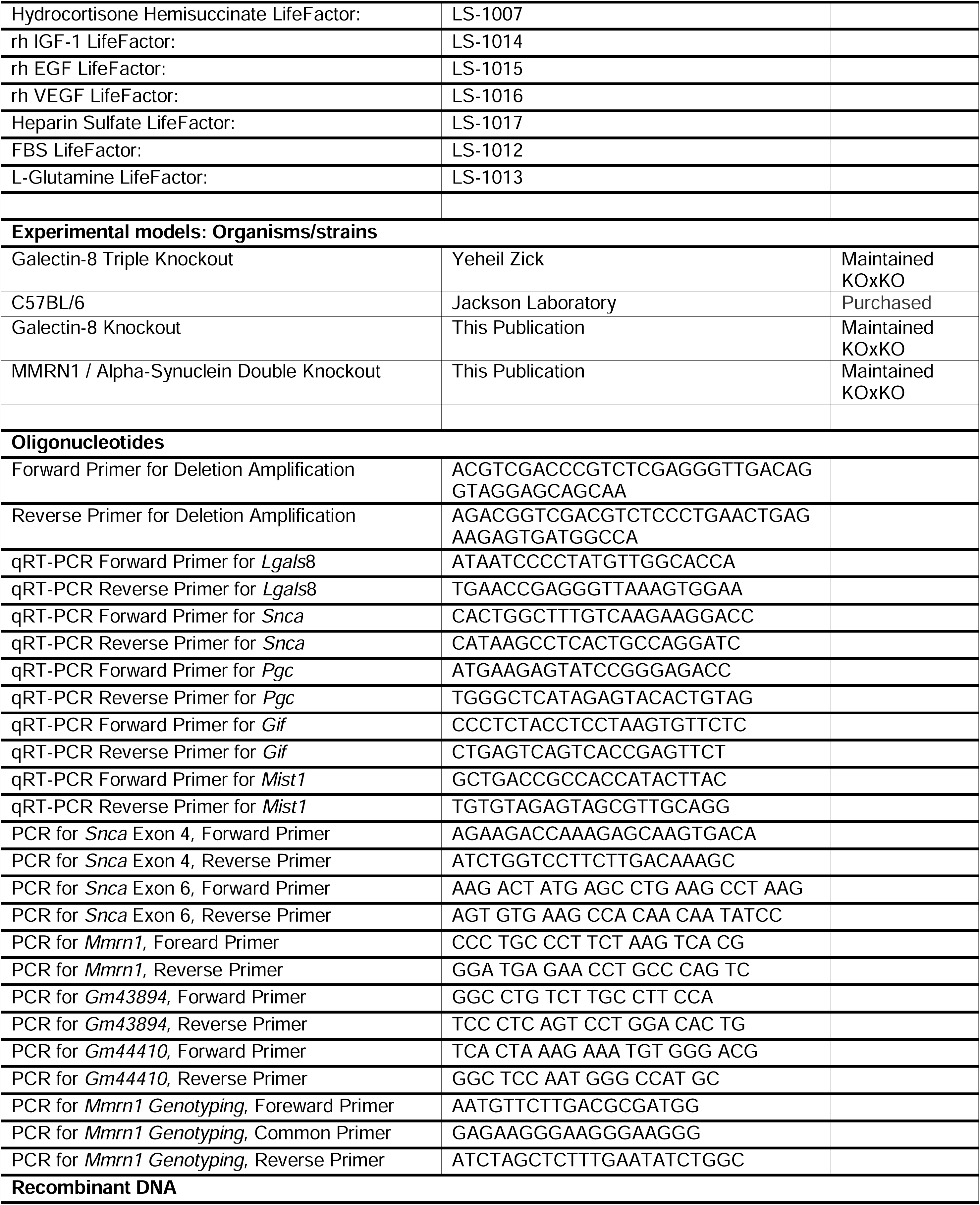
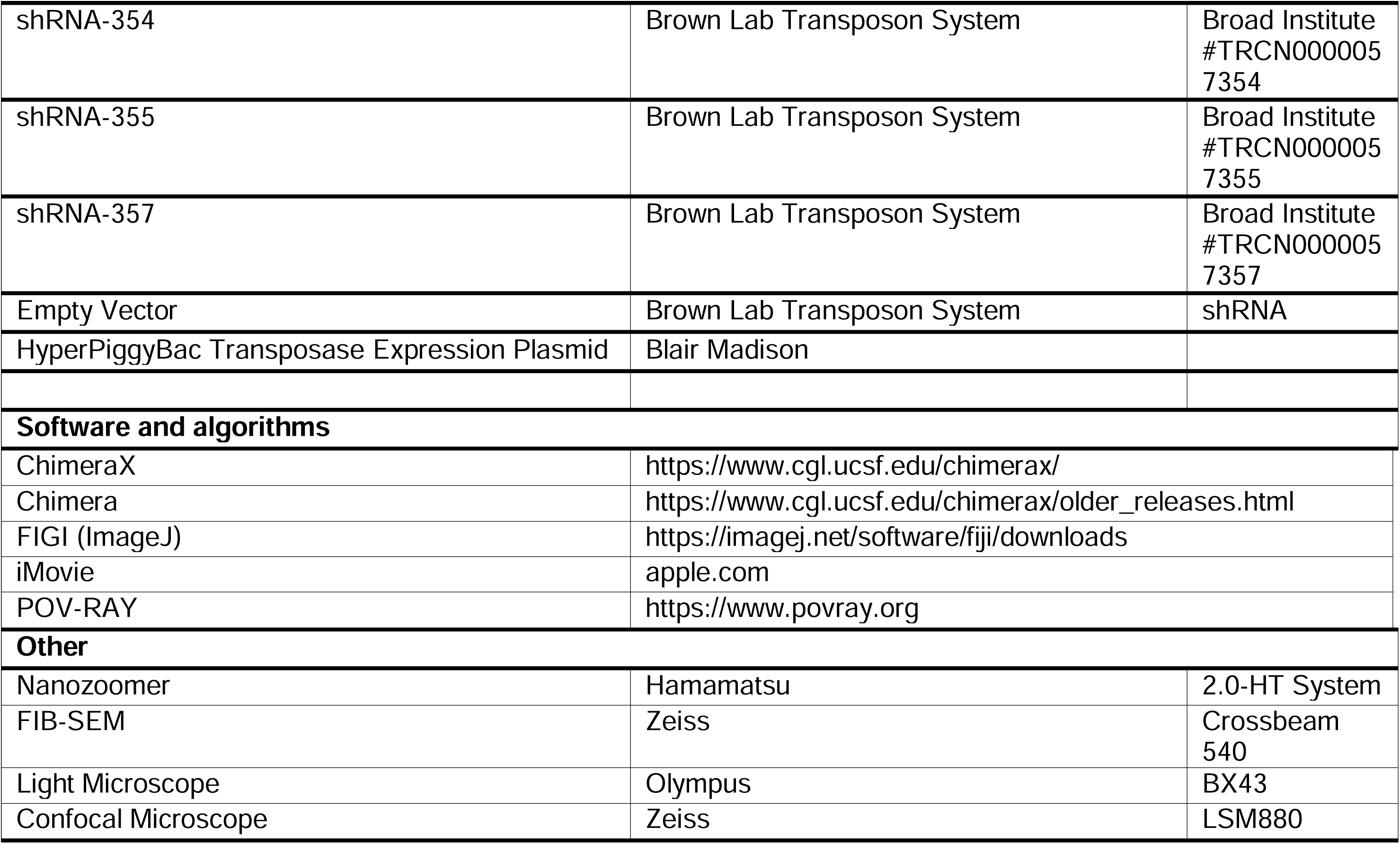

## Abbreviations used in this paper

SPEM: Spasmolytic Polypeptide Expressing Metaplasia
Pan-IN: pancreatic intraepithelial neoplasia
CRD: Carbohydrate Recognition Domain
RNA: Ribonucleic acid
RNAseq: RNA sequencing
scRNAseq: single cell RNA sequencing
qRT-PCR: quantitative real-time Polymerase Chain Reaction
shRNA: short hairpin RNA
TGF: Transforming Growth Factor
WNT: Wingless-related integration site

## ACKNOWLEDGEMENTS

We thank Linda Samuelson for very helpful and thoughtful discussion and advice. The work was supported by the following grants: **<u>JWB</u>**: K08 DK132496; Department of Defense W81XWH-20-1-0630, R21 AI156236, P30 DK052574 and the Foundation for Barnes-Jewish Hospital**. <u>JBS</u>:** R01 DK141682, Foundation for Barnes-Jewish Hospital.

## RESOURCE AVAILABILITY

### Lead Contact

Further information and request for resources and reagents should be directed to and will be fulfilled by Lead Contact, Jeffrey W. Brown

### Materials Availability

All mouse lines and materials used in this study were provided or purchased from the mentioned companies or researchers. This study did not generate any new or unique reagents. If the mouse lines are still in the investigator’s colony, we are happy to share them.

### Data and Code Availability

**-**Data is available on request.

-The paper does not report original code.

-Any additional information or data is available from the lead contact upon request.

---

**Supplemental Figure 1.**
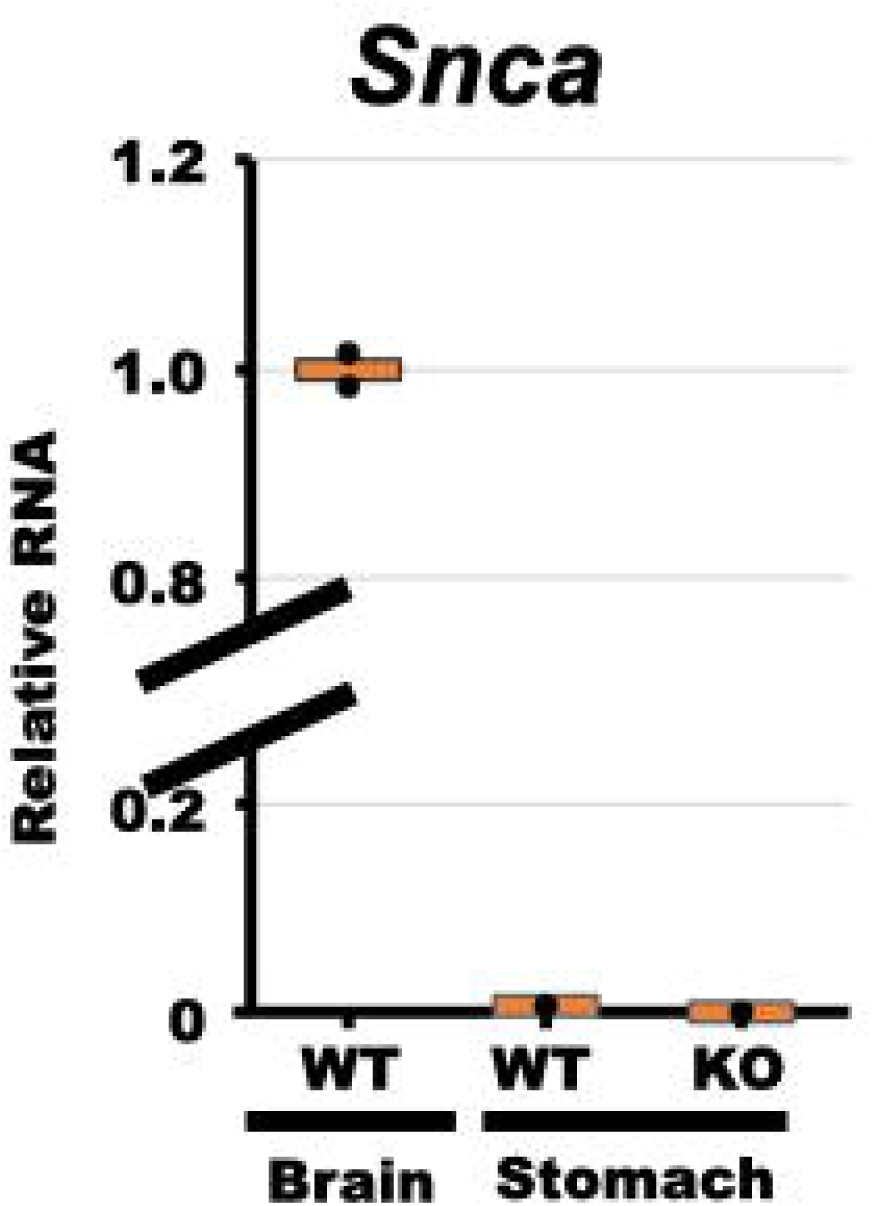
Alpha-Synuclein RNA is not detected in the mouse stomach at homeostasis. mRNA determined by qRT-PCR of alpha-synuclein from mouse brain (positive control), compared to stomach from wild-type C57BL/6J and Mmrn1^KO^/Scna^KO^ at 6 weeks of age.

**Supplemental Table 1.**
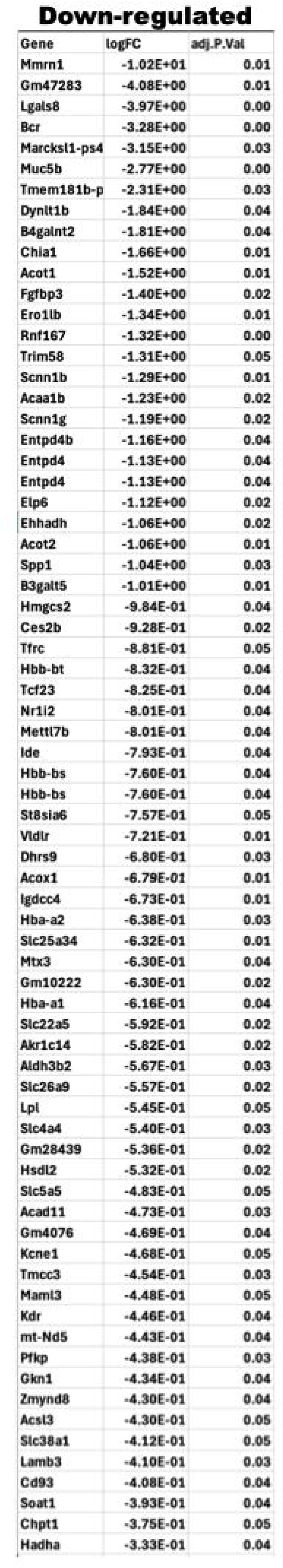
List of Genes Significantly Downregulated in *Lgals8^−/−^/Mmrn1^−/−^ /Snca^−/−^* relative to Wild-Type C57BL/6J.

**Supplemental Table 2.**
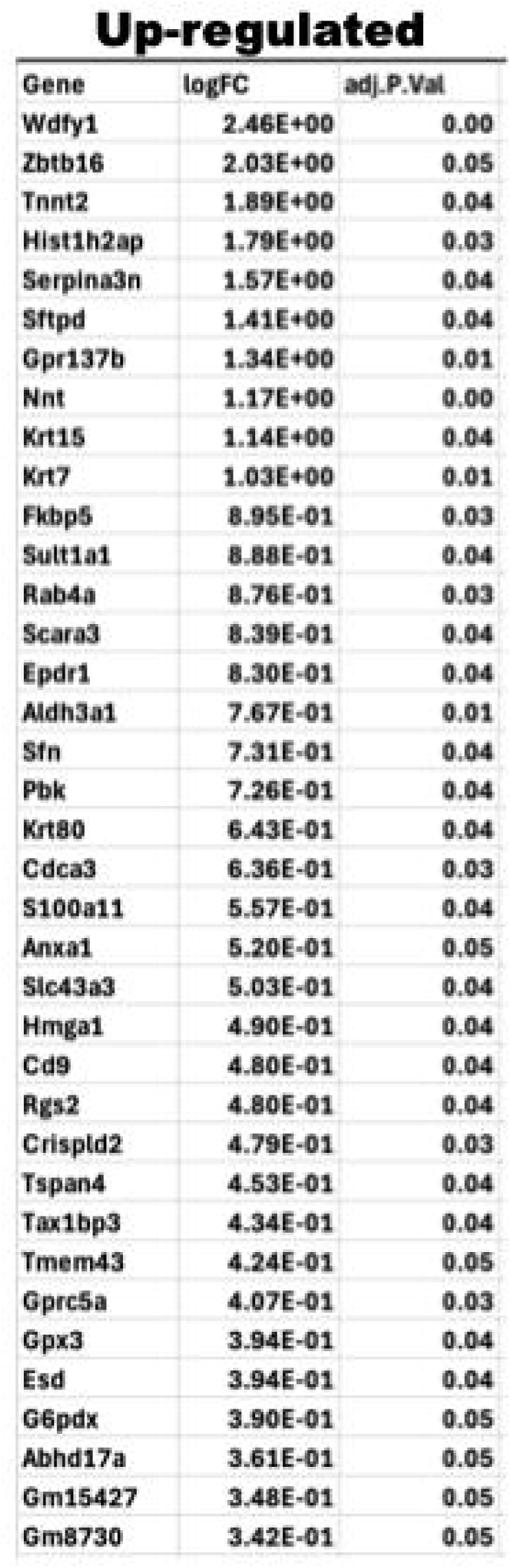
List of Genes Significantly Upregulated in *Lgals8^−/−^/Mmrn1^−/−^/Snca^−/−^* relative to Wild-Type C57BL/6J.

